# A novel technique for monitoring Alzheimer’s disease associated changes in brain-derived extracellular vesicle cargos in mouse models

**DOI:** 10.64898/2026.03.13.711599

**Authors:** Nicholas Francis Fitz, Md Shahnur Alam, Mary Ann Ostach, Sia Garg, Iliya Lefterov, Radosveta Koldamova

## Abstract

Extracellular vesicles (EVs) are critical mediators of intercellular communication, carrying molecular cargos such as small noncoding RNAs (ncRNAs) that reflect the physiological and pathological state of their cells of origin. However, studying brain-derived EVs has been challenging due to the blood-brain barrier. Here, we optimized and validated an open-flow microdialysis (OFM) protocol for sampling EVs directly from brain interstitial fluid (ISF) in wild-type and APP/PS1 transgenic mice. *Ex-vivo* validation using plasma EVs demonstrated that OFM effectively captures the full EV population. *In-vivo* cerebral OFM (cOFM) enabled successful collection of brain ISF EVs, which were characterized by nanoparticle tracking analysis (NTA), electron microscopy, and western blotting, confirming their similarity to EVs isolated directly from brain tissue and plasma. Identification of small ncRNA cargos revealed that EVs sampled from brain ISF by cOFM were enriched in brain-specific signatures, many of which are associated with neuronal cell populations and biological functions. Furthermore, we observed a unique small ncRNA signature from the brain ISF EVs in the Alzheimer’s disease preclinical model compared to wild-type mice. These small ncRNAs were associated with genes considered important in biological functions associated with neurodegeneration. Our findings demonstrate that cOFM is a powerful tool for *in-vivo* sampling of brain EVs and highlight the unique molecular landscape of ISF EV small ncRNA cargos. This study offers new opportunities for biomarker discovery and mechanistic insights into neurodegenerative diseases, such as Alzheimer’s disease.

## 1. Introduction

Extracellular vesicles (EVs) are lipid membrane-delimited nanoscale vesicles secreted by almost all cell types into the extracellular space. EVs are heterogeneous in size and classically described as exosomes (30-100nm), microvesicles (100nm-1µm) and apoptotic bodies (1-5 µm) (Hadizadeh, Bagheri et al. 2022), each produced by distinct cellular processes and containing unique cargos. The 2023 guidelines for studies on extracellular vesicles (Minimal Information for Studies of Extracellular Vesicles 2023; MISEV2023) recommends the general term “EV” as the most appropriate to describe small (30-150 nm), large (100-1000 nm), and very large (1-10 µm) EVs released by cells (Welsh, Goberdhan et al. 2024). For the last two decades there has been increased research to better understand the biological role of small EVs. Biogenesis of small EVs is initiated by the invagination of the cellular plasma membrane and formation of early endosomes, which mature into multivesicular bodies (MVBs). The MVB can be directed to combine with lysosomes for degradation or formation of EVs through fusion with the plasma membrane, a process dependent on the endosomal sorting complex that is required for transportation (ESCRT) (Hadizadeh, Bagheri et al. 2022). Initially, EVs were simply perceived as released recycling bins (Harding, Heuser and Stahl 1983), exporting byproducts and cellular toxins; now their importance in intercellular communication is accepted as they carry bioactive cargos including: non-secreted proteins, nucleic acids, lipids and metabolites (Witwer and Thery 2019). Notably small EVs, regardless of their origin, are enriched with extracellular RNAs of many different types, particularly small noncoding RNAs (ncRNAs). The most frequently reported types of small ncRNAs carried by small EVs include: miRNAs (microRNAs), tRNAs (transfer RNAs), piRNAs (PIWI-interacting RNAs), snRNAs (small nuclear RNAs), snoRNA (small nucleolar RNAs), and circRNAs (circular RNAs). While the exact functions of many small ncRNAs are not fully known, studies show their ability to regulate gene expression at the level of messenger RNA (mRNA) processing (Telonis, Loher et al. 2015, Becker, Hirsch et al. 2019, Gebert and MacRae 2019). For example, miRNAs bind to a specific sequence of target mRNAs to induce translation repression through either mRNA silencing or degradation. First it was shown that miRNAs bind the 3’ untranslated region (UTR) but more recently binding sites have been identified in the 5′ UTR, coding sequence, and gene promoter regions (O’Brien, Hayder et al. 2018). In contrast, circRNAs can act as microRNA sponges, inhibiting miRNA modulation of gene expression; or bind to the 3′ UTR of mRNAs to induce their decay (Li, Wang and Jin 2021). Thus, EVs carry small ncRNAs which can act either as activators or suppressors of gene expression in a complex relationship (Whiteside 2016, Barros, Carneiro et al. 2018).

In brain EVs are critical mediators of intercellular communication, enabling transfer of molecular signals between neurons and glial cells and contributing to brain development and neurodegenerative pathological processes (Lizarraga-Valderrama and Sheridan 2021). Neuron-derived small EVs can influence postsynaptic signaling and support neural circuit formation during development (Korkut, Li et al. 2013). Emerging evidence also implicates small EV cargos, such as small RNAs, in regulating neuron-glia interactions and brain homeostasis. For instance, neuronal mir-21-5p released via small EVs has been shown to modulate microglial activation, astrocyte gene expression, and coordinate a pro-inflammatory response (Ge, Han et al. 2015) while mir-615-5p in microglia-derived EVs containing impairs myelin regeneration by inhibiting oligodendrocyte precursor differentiation (Ji, Guo et al. 2024). Collectively, these findings support the role of small EVs as mediators of cell-to-cell communication in brain, impacting brain development, homeostasis, neuroinflammation, and neurodegeneration.

EVs have been at the center of research towards the identification of molecular biomarkers for diagnosis, prognosis and assessment of therapies in numerous diseases, including Alzheimer’s disease (AD) (Gomes, Tzouanou et al. 2022). For example, studies have shown that EVs isolated from AD patients and mouse models contain hyperphosphorylated tau, which is believed to be important in tau spreading (Winston, Aulston et al. 2019, Ruan, Pathak et al. 2021). Microglial-derived EV cargos, such as mir-711, alleviates neurodegeneration and cognitive deficits in AD mouse models (Zhang, Xu et al. 2020). Additionally, mir-873a-5p cargo from astrocyte-derived EVs reduces neuroinflammation by suppressing NF-κB signaling (Long, Yao et al. 2020). We were the first to show differences in the small ncRNA profiles of plasma EVs isolated from AD patients compared to health controls, specifically demonstrating increases in the abundance of brain specific SNORD115 and SNORD116 in AD which showed significant discriminatory power as a biomarker (Fitz, Wang et al. 2021). Therefore, EV cargos, including small ncRNAs, can play a pivotal role in pathophysiologic processes that promote the development and progression of neurodegeneration and represent novel disease biomarkers.

Recently, EVs have been isolated from plasma (Fitz, Wang et al. 2021, Badhwar, Hirschberg et al. 2023, Lucien, Gustafson et al. 2023), CSF (Thompson, Gray et al. 2020, Guha, Misra et al. 2023), postmortem brain tissue from AD patients (Vella, Scicluna et al. 2017, Huang, Arab et al. 2023) and AD mouse models (Polanco, Scicluna et al. 2016, Bodart-Santos, Pinheiro et al. 2023). However, each compartment yields disparate results and has unique limitations. While CSF provides a direct window into brain EV populations there are limits to the amount of CSF which can be collected from AD mouse models which greatly limit the concentration of EVs for characterizing cargos (Sandau, Magana et al. 2024). In contrast, brain tissue offers a direct source of brain derived EVs but requires invasive biopsies or post-mortem samples, and tissue EV isolation techniques utilize mechanical and enzymatic dissociation steps that risk vesicle disruption, cargo modifications and contamination with intracellular components, potentially confounding downstream analyses (Vella, Scicluna et al. 2017, Hurwitz, Sun et al. 2018). Characterizing brain-derived EVs from plasma also has limitations, as they make up only a small portion of all circulating EVs and immunoaffinity approaches used to isolate brain-derived EVs have come under scrutiny due to non-specific binding, contamination by soluble proteins, limited marker specificity, and potential loss of diverse EV subpopulations (Norman, Ter-Ovanesyan et al. 2021, Kadam, Wacker et al. 2025). Even with the advancement of protocols for isolating EVs most of these studies are designed to test a single time point and could potentially miss dynamic changes in EV cargos associated with brain developmental windows and stages of neurodegeneration. Therefore, single timepoint analysis of EVs may not fully explain the active role of EVs and potentially bias understanding of their importance during development or disease progression.

To fill the gap in research targeting brain derived EVs and to gain a more comprehensive understanding of their role, we developed a novel *in vivo* microdialysis technique which allows for real time collection from brain interstitial fluid (ISF) where EVs are directly released. We applied cerebral Open Flow Microdialysis (cOFM) method for collecting EVs from the brain ISF of awake freely moving mice and subsequently isolated EVs from ISF, plasma and brain tissues of wild-type (WT) and APP/PS1 mice for compartment specific characterization of small EV cargo, especially small ncRNAs. We identified a significant number of differentially enriched small ncRNAs transcripts when comparing EVs isolated from ISF and plasma, suggesting isolation of unique EV populations associated with the brain utilizing the cOFM technique. Further, we determined unique small ncRNAs profiles of EVs from the ISF in APP/PS1 mice compared to WT controls. Thus, the novel cOFM technique for sampling EVs from the ISF will be an important method to better understand changes in EV cargos associated with brain development, different neurodegenerative diseases, and therapeutic monitoring.

## 2. Materials and Methods

### 2.1 Animals

All animal studies were approved by the University of Pittsburgh Institutional Animal Care and Use Committee, which adheres to the guidelines outlined in the Guide for the Care and Use of Laboratory Animals from the United States Department of Health and Human Services. Six to nine-months old APPswe, PSEN1dE9 transgenic mice [B6.Cg-Tg (APPswe, PSEN1ΔE9)85Dbo/J] (APP/PS1) or wild-type (WT) littermate controls with equal male to female ratio were used for this study. Mice were initially purchased from The Jackson Laboratory (USA); and animals for experimental use were bred in-house and used experimentally as heterozygous along with WT littermates. All experimental mice were kept on a 12 h light-dark cycle with ad libitum access to food and water.

### 2.2 *Ex vivo* validation of EV sampling with open flow microdialysis

Plasma samples were collected from 4 WT mice (2 males and 2 females) and samples were then centrifuged for 15 min at 3000 g, 4°C. The supernatant was transferred to a new test tube and centrifuged for 15 min at 3000 g, 4°C. We used an Automatic Fraction Collector (AFC) with a qEV Original Column, (SP5, IZON) to isolate plasma EVs. First, the qEV Original Column was flushed with 15 mL of particle free 0.1 M PBS, pH 7.4 and 500 µL plasma was transferred to the column. 9 mL of particle free 0.1 M PBS, pH 7.4 was added to column when prompted and 10 fractions of 0.5 mL volume for each plasma sample were collected. Fractions 7 through 10 were combined and centrifuged for 1 hour at 34,600 RPM, 4°C with no brake (Beckman Optima LE80K ultracentrifuge). 1.5 mL of the supernatant was removed and the remaining 500 µL of was used to resuspend the EV pellet. Half of the isolated plasma EVs were used for the *ex vivo* Open Flow Microdialysis (OFM) experiment and the remaining EV sample was aliquoted for characterization. In brief, a perfusate bag (BASi) filled with 0.1 M PBS, pH 7.4 and tubing was attached to the peristaltic pump (MPP 102 PC, BASi) and the guide canula with sampling insert was lowered into 160 µL of isolated plasma EV samples. We sampled the *ex vivo* plasma EV samples at 1 μL/min until 160 µL of perfusate was collected for each sample on ice. EVs sampled with the OFM were isolated from the perfusate and characterized as described below.

### 2.3 Surgical implantation of guide cannulas and cerebral OFM sampling of brain interstitial fluid

cOFM guide cannula implantation was performed according to previously published methods (Fitz, Cronican et al. 2013, Shahnur, Nakano et al. 2021). Following induction of anesthesia with 5% isoflurane, the mouse was moved to the stereotaxic frame, where the head was secured, core body temperature maintained at 37 °C using a heating pad and anesthesia continued with 2% isoflurane. The head was shaven, surgical site sterilized with two separate iodine-alcohol washes, a 50% mixture of bupivacaine and lidocaine applied to the surgical site and ophthalmic ointment applied to the eyes. The scalp was spread with a midline incision exposing the dorsal aspect of the skull and leveled. A 4.5 mm diameter craniotomy was performed over the left parietal cortex using a dental drill. A cOFM guide cannula (cOFM-GD-6-2, BASi) was implanted at the target coordinate (A/P: -2.0 mm, M/L: - 1.5 mm, D/V: -3.5). The cOFM probe was fixed using two bone screws and binary dental cement. The surgical site was sutured, triple antibiotic cream applied, Buprenex (0.1 mg/kg; IP) provided for analgesia and sterile saline administered for rehydration. Mice were allowed to recover on a heating pad, until freely mobile, before returning to their home cage. After two weeks recovery period for reestablishing the brain barriers, the cOFM sampling procedure was started. Each mouse was shortly anaesthetized using isoflurane to replace the healing dummy cannula with the sampling insert (cOFM-S-8, BASi). The sampling insert was connected to the inlet and outlet tubing via tubing connectors and both tubings were operated by a peristaltic pump (MPP 102 PC, BASi). Prior to insertion into the guide cannula the tubing was flushed at 5 μL/min for 5 min with commercially available artificial cerebrospinal fluid (597316, Harvard Apparatus) and checked for leaks or air bubbles. Subsequently, the flow rate was lowered to the sampling flow rate of 1 μL/min and samples were collected in a refrigerated fraction collector (CMA 470). Throughout the cOFM collection, the animals were freely moving and had access to food and water.

### 2.4 EV isolation from brain interstitial fluid, plasma and brain

EVs from plasma and cOFM were isolated using Plasma/Serum Exosome Purification Mini Kit (57400, Norgen Biotech Corp.) according to the manufacturer instruction. 400 µL plasma was used for EV isolation in which 3.6 mL particle free water was added to obtain a 4 ml volume equal to that collected with the cOFM. 100 µL of ExoC Buffer was added to each 4 mL sample followed by 200 µL of Slurry E and incubated at room temperature for 5 min. Following incubation, the samples were centrifuged for 2 min at 2,000 RPM, the supernatant was discarded and 200 µL ExoR Buffer added to the slurry pellet for releasing the EVs. After incubation for 5 min, samples were centrifuged for 2 min at 500 RPM and supernatant transferred to a mini filter spin column assembly and centrifuged for 1 min at 6000 RPM to capture the EVs in the flowthrough.

EVs were isolated from fresh frozen brain (half hemisphere excluding subcortical and cerebellum regions) with minor modification of the following protocol (Huang, Cheng et al. 2020). First, the brain was sliced with a razor blade into small sections 3 x 3 mm on dry ice, and then dissociated in Hibernate-E medium (A1247601, Thermo Fisher Scientific) containing 75 U/mL collagenase type 3 (CLS-3, S8P18814; Worthington Biochem Corp.) at 37°C for 15 min. After incubation, PhosSTOP (04906837001, Millipore Sigma) and Complete Protease Inhibitor (11697498001, Millipore Sigma) were added to terminate digestion and sample volume normalized to 4 ml with PBS. The samples were sequentially centrifuged at 300 × g for 10 min, and the supernatant at 2000 × g for 15 min at 4°C. This supernatant was then centrifuged at 7500 RPM (SW 40Ti rotor, Beckman Optima LE80K ultracentrifuge) for 30 min at 4°C. The supernatant was concentrated with a 100 kilodalton (kDa) MWCO protein concentrator (88524, Thermo Fisher Scientific) from 10 ml to 500 µl. The 500 µl sample was separated on a size-exclusion chromatography column (qEV 70 nm, IZON Science) as above with collection of fractions 7-10 followed by concentration with 10 kilodalton (kDa) MWCO protein concentrator (88517, Thermo Fisher Scientific) to a final volume of 200 µl.

### 2.5 EV characterization

For EV characterization, 1 ml of diluted EV sample was loaded on a NS300 Nanosight instrument (Malvern Instruments) at a rate of 400 μL/min. The instrument was calibrated with 100 nm latex beads diluted in particle free PBS prior to running any set of samples. Before and after the analysis of each sample, particle free PBS was used to wash the instrument. Sample analysis consisted of 3 separate measurements of 30 seconds each. Nanoparticle Tracking Analysis (NTA) software, version 3.4, was used to visualize and measure each measurement. EV samples were further characterized by Western blotting and Transmission electron microscopy (TEM). For Western blotting, EV proteins were resolved on 4–12% Bis-Tris gels (NW04122, Thermo Fisher Scientific) and transferred onto nitrocellulose membranes (IB23001, Thermo Fisher Scientific, iBlot2 Gel system). These membranes were probed with anti-Calnexin (abc22595, 1:2000 dilution, secondary antibody-Santa Cruz goat anti-rabbit IgG-HRP, 1:3000); anti-Flotilin (BD Biosciences 610820, 1:1000 dilution; secondary antibody-Santa Cruz goat anti-mouse IgG-HRP, 1:3000); anti-CD63 (System Biosciences EXOAB-CD63A-1, 1:500 dilution, secondary antibody-goat anti-rabbit HRP, 1:10,000) and anti-CD81 (System Biosciences EXOAB-CD81A-1, 1:500 dilution, secondary antibody-goat anti-rabbit HRP, 1:10,000). Immunoreactive signals were visualized using enhanced chemiluminescence with the Amersham Imager 600 (GE Lifescience). TEM was performed on freshly isolated EVs from ISF as previously described (Fitz, Wang et al. 2021). Briefly, EVs were adhered to copper grids coated with 0.125% Formvar in chloroform, stained with 1% uranyl acetate in ddH₂O, and imaged immediately using a JEM 1011 transmission electron microscope.

### 2.6 RNA isolation and library preparation

Total RNA was extracted from isolated EVs using Exosomal RNA Isolation Kit (58000, Norgen Biotech Corp.) according to the manufacturer’s protocol. The quantification and quality of isolated RNA was evaluated with the Agilent 2100 Bioanalyzer RNA Pico assay (50671513, Agilent technologies) following the manufacturer’s instructions. RNA was pelleted with Pellet Paint (690493, Novagen) and reconstituted in 6 µl of nuclease free water. The NEBNext small RNA sample library preparation kit (E7300S, New England Biolabs) was used according to manufacturer’s protocol with minor modifications. All reactions were with a 1:10 adapter dilution and run with 18 PCR cycles. The library products were cleaned using AMPure XP beads (A63881, Beckman Coulter). The small RNA library quality and quantity were assessed on Bioanalyzer 2100 High Sensitivity DNA chip (50674626, Agilent technologies) and sequenced at the Health Sciences Sequencing Core at UPMC Children’s Hospital of Pittsburgh, on the NextSeq 2000 machine (1×51 bp).

### 2.7 Computational analysis of sequencing data

Small noncoding RNA library data generated from all compartments and from published data of the mouse tissue atlas (Isakova, Fehlmann et al. 2020) were aligned and quality checked by STAR (v 2.5.3a) and annotated with COMPSRA (v 1.0.3) and differential abundance determined by DESeq2 (v 1.36.0). Predicted miRNA gene targets were analyzed using TargetScanMouse 8.0 at cumulative weighted context score cutoff<-0.5, while circRNA, associated genes were identified by mouse circBase database (www.circbase.org). Functional annotation clustering was performed using the Database for Annotation, Visualization and Integrated Discovery (DAVID) with all GO terms considered significant at *p* < 0.05. Furthermore, GO enrichment analysis of predicted target genes was performed using Metascape (http://metascape.org), a free web-based platform for integrative gene function analysis. Hypergeometric distribution testing for commonly expressed genes between comparisons was performed using phyper in an R environment.

### 2.8 Statistical Analysis

Sample size information for experiment is included in the figure legends with each sampling of males and females. EV characterization was analyzed with a two-tailed unpaired t-test (GraphPad Prism v10.0.3). Biological function terms were reduced and visualized utilizing REVIGO, reduce+visualize Gene Ontology, (v 1.8.1).

## 3 Results

### 3.1 Validation of Open-Flow Microdialysis (OFM) for EV sampling

Microdialysis has been useful in dynamically assessing the concentration of a variety of neurotransmitters and secreted proteins in brain ISF. Given the presence of blood brain barriers, neural release of EVs is likely to produce a unique population of EVs whose cargos can change during brain development and neurodegeneration. We wanted to develop an optimized protocol for sampling of EVs from brain ISF utilizing OFM. We began validation of the utility of OFM for EV sampling *ex vivo* utilizing 4 plasma EV samples collected from WT mice. EVs were isolated from plasma as described in the methods and we assessed the mean size and concentration of EVs prior and after sampling with OFM with nanoparticle tracking analysis (NTA). EVs from the OFM samples showed similar size distribution of those isolated directly from plasma indicating that the OFM protocol was sampling the entire population of EVs **(Figure 1A**). OFM sampled EVs had an average concentration of 2.36 × 10⁸ particles/mL and a mean particle size of 148 nm; which is characteristic of small EVs and closely matched the characterization of EVs directly isolated from plasma (average concentration 2.07 × 10⁸ particles/mL and mean particle size 142 nm) (**Figure 1B-C**). Western blot analysis confirmed the presence of canonical EV markers, including CD63, CD81, and flotillin-1 in the OFM sampled EVs and at similar levels as EVs directly isolated from plasma, supporting the purity and identity of the small EVs sampled with the OFM protocol (**Figure 1D**). These findings demonstrate that OFM is a viable and highly efficient method for capturing EVs *ex vivo* and has the potential for EV sampling from tissue microenvironments for downstream molecular EV cargo analysis.

**Figure 1.**
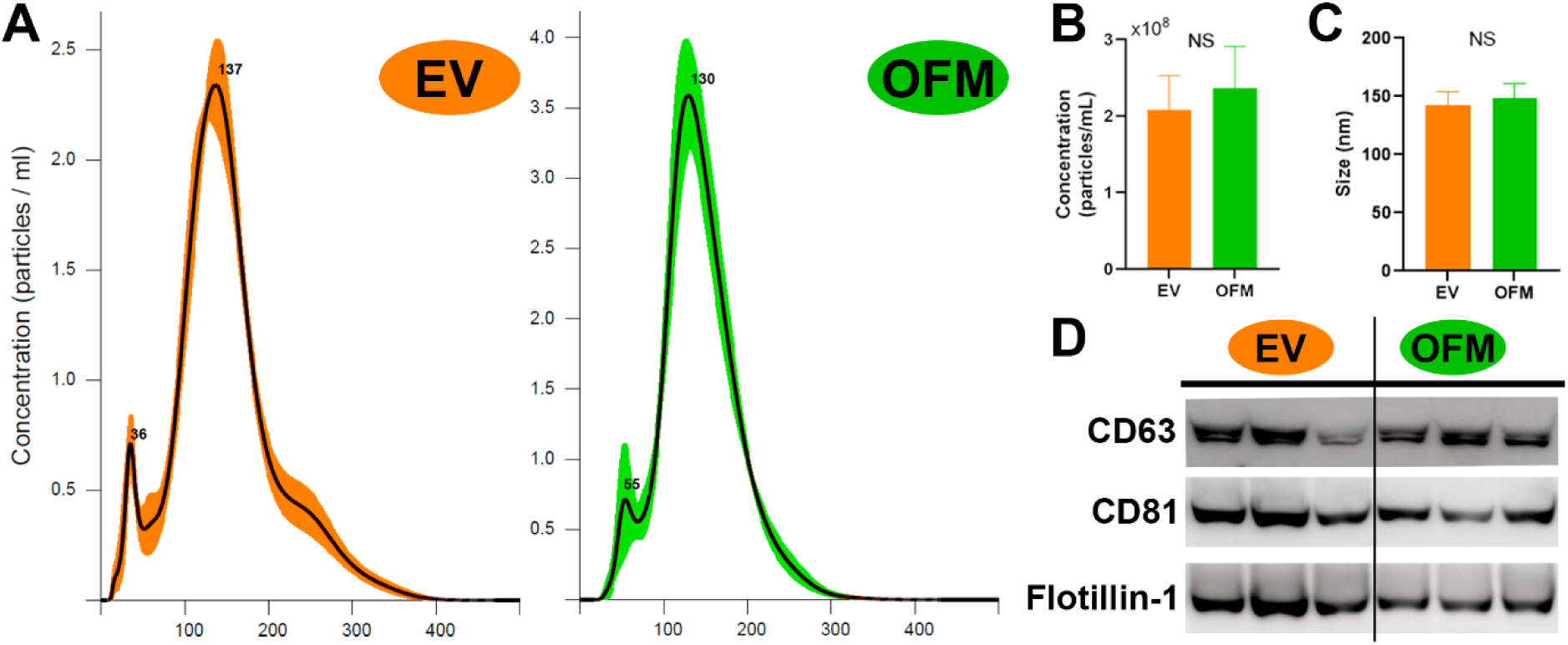
*Ex-vivo* validation of extracellular vesicle (EV) recovery utilizing OFM. To validate the recovery of EVs we performed OFM on plasma EVs *ex-vivo* and compared them with aliquots of the same sample of directly isolated plasma EVs. (**A**) EVs particle size distribution profiles measured by nanoparticle tracking analysis (NTA) of plasma EVs and OFM EVs shows similar distribution of EV size following OFM sampling. Bar graphs representing the particle concentration (**B**) and mean size (**C**) were not significantly different between plasma EV or OFM EV samples. (**D**) Western blots of EV markers CD63, CD81 and Flotilin-1 show similar band intensity between plasma EV or OFM EV samples. Bar graphs represent mean ± SEM; n=4 WT mice samples. Statistical significance was determined by unpaired t-test. NS, not significant.

### 3.2 Efficient *in vivo* sampling of EVs from mouse brain interstitial fluid utilizing cerebral open flow microdialysis (cOFM)

After *ex vivo* validation of EV sampling using OFM, we initiated an *in vivo* experiment for sampling EVs from brain ISF utilizing the cOFM technique in WT and APP/PS1 mice. Following collection of the cOFM perfusate of the ISF, plasma and brain were collected from all animals. EVs were then isolated from all three compartments (ISF, plasma, brain) using our described EV isolation protocols. We first characterized the EVs sampled from the ISF with cOFM based on size, concentration and protein composition and compared them to EVs isolated from plasma and brain. NTA showed similar distribution of EV size from brain tissue (**Figure 2A**), plasma (**Figure 2B**) and ISF (**Figure 2C**). Transmission electron microscopy (TEM) further confirmed the presence of intact EVs isolated from the ISF, with the typical round morphology and ∼100 nm size (**Figure 2D**). NTA measured an average EV concentration of 2.81 x 10^10^ particles/mL in brain, 1.31 x 10^10^ particles/mL in plasma, and 2.07 x 10^8^ particles/mL in ISF (**Figure 2E**). There was no significant difference in the EV size from the three compartments with an average of 180.6 nm in brain, 133.1 nm in plasma, and 155.7 nm in ISF (**Figure 2F**). The predominance of particles measured between 50 nm and 200 nm which indicates an enrichment of small EVs. Finally, the enrichment of small EVs was confirmed by western blotting using canonical EVs markers flotillin-1 and CD81 and cellular debris marker calnexin (**Figure 2G-I**). We observed enrichment of these markers from the EVs isolated from all three compartments with a lack of calnexin, suggesting pure EV isolations from brain, plasma and ISF with minimal cellular debris. These findings demonstrate successful isolation and characterization of EV sampled from the ISF with the cOFM procedure which were of similar size and concentration as those isolated from plasma and brain. This sampling of EVs from ISF utilizing cOFM technology supports downstream comparative analyses and suggests the potential for identifying compartment specific EV cargo signatures. Furthermore, this highlights the utility of cOFM for sampling EVs from the ISF of different mouse models utilized for assessing brain development and neurodegeneration.

**Figure 2.**
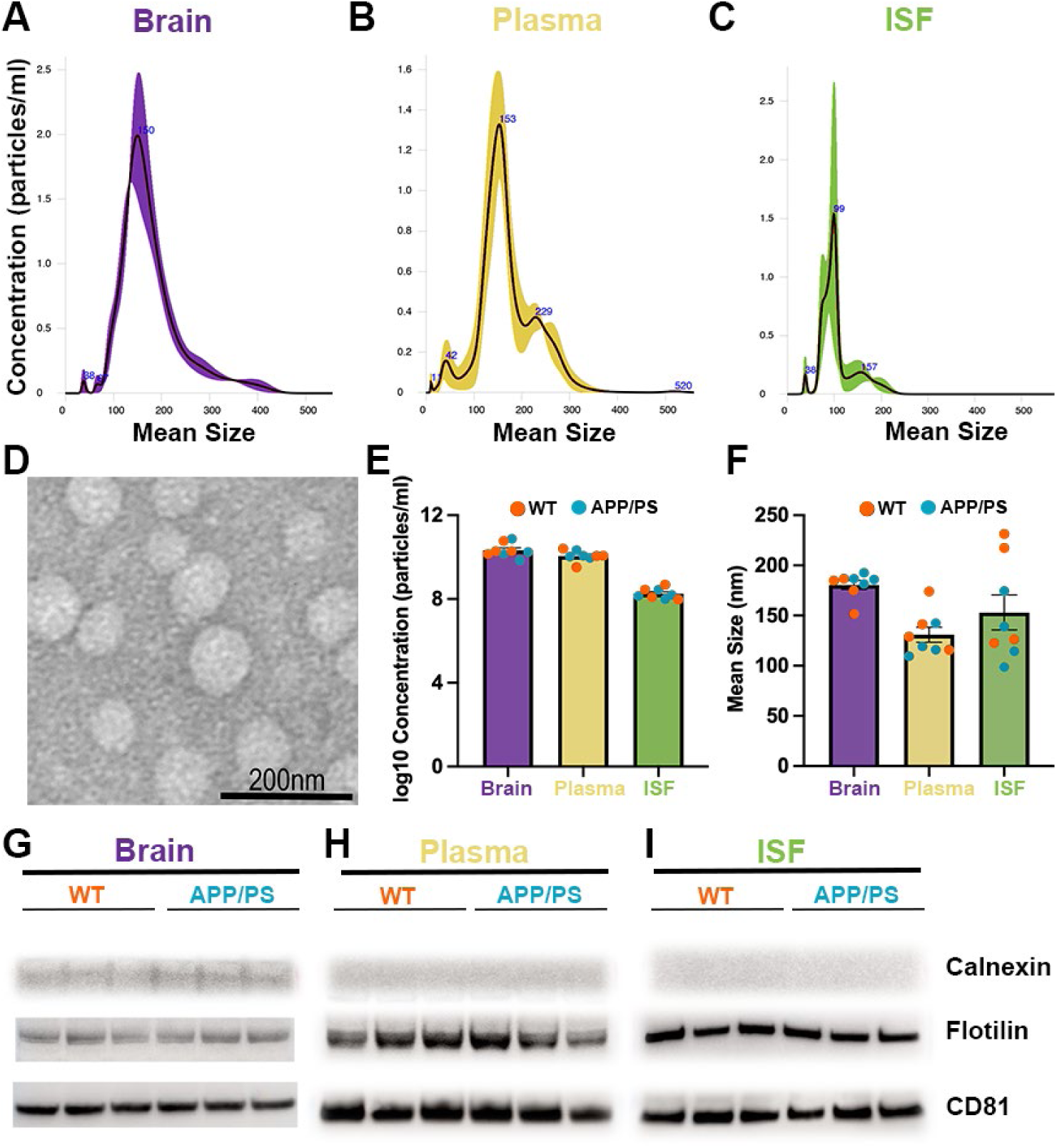
Characterization of EVs sampled from brain ISF by cOFM and isolated from brain tissue and plasma. EVs particle size distribution profiles measured by NTA of brain EVs (**A**), plasma EVs (**B**) and cOFM sampled ISF EVs (**C**) showing similar size distribution from the three compartments. (**D**) Representative TEM image to visualize size and morphology of EVs sampled from brain ISF by cOFM. Bar graphs representing EV concentration (**E,** log10 transformed) and mean size (**F**) as measured by NTA analysis for EV isolated from brain, plasma and ISF. EVs from all three compartments show similar mean size and ISF EVs had slightly lower concentration as compared with EVs isolated from either plasma or brain. There was no significant difference in EV concentration or size from all three compartments between APP/PS1 and WT mice. Representative western blot images of EV markers Flotilin-1 and CD81 and negative marker Calnexin for brain (**G**), plasma (**H**) and cOFM (**I**). Bar graphs represent mean ± SEM; n = 8 mice/group (4 WT and 4 APP/PS1 with equal sex distribution).

### 3.3 Unique small ncRNA signatures were associated with EVs isolated from ISF compared to plasma

Following the initial characterization of EVs from the three compartments (ISF, plasma and brain), we wanted to determine if EVs sampled from the ISF are a unique population of EVs compared to those isolated from the plasma. EVs from the ISF are directly shed from brain-resident cells, including neurons, astrocytes, and microglia, thereby providing a real-time window of the molecular changes occurring within the central nervous system and offer high specificity whose access is limited. In contrast, plasma EVs provide a minimally invasive avenue to capture systemic and brain-derived signals, reflecting peripheral surrogates of central nervous system changes. In this study, we focus on EVs isolated from ISF and plasma to better understand the differences between brain-specific and peripheral small ncRNA signatures associated with these EVs. We isolated total RNA from ISF and plasma EVs and generated small ncRNA libraries to determine differences in the abundance of small ncRNA cargos from 4 WT and 4 APP/PS1 mice. We mapped and annotated the expression of distinct small ncRNA classes: circRNA, snoRNA, snRNA, miRNA, and tRNA from ISF and plasma EVs using COMPSRA (Li, Kho et al. 2020). We identified a total of 12,104 distinct circRNA, 123 snoRNA, 496 snRNA, 648 miRNA, and 439 tRNA transcripts in plasma EVs which correspond to 73.6%, 3%, 12%, 20%, and 93% of annotated transcripts of each respective class. Similarly, in EVs sampled from the ISF we identified a total of 12,394 distinct circRNA, 124 snoRNA, 239 snRNA, 603 miRNA, and 444 tRNA transcripts in plasma which correspond to 75%, 3%, 14%, 18.6%, and 94% of annotated transcripts of each respective class. These findings indicate a high degree of overlap and minimal variation in the numbers of small ncRNAs annotated when comparing EVs isolated from ISF and plasma (**Figure 3A**). To determine the brain-enrichment of our ISF sampled EVs, we first compared the number of small ncRNAs expressed in our dataset with that of Isakova et al. which reported a mouse tissue-specific atlas of small ncRNAs, identifying enriched small ncRNAs in brain and other tissues (Isakova, Fehlmann et al. 2020). We observed that many miRNAs identified in our EVs sampled from brain ISF (279 species) were also enriched in brain tissue from the tissue atlas. By chance, this was greater than predicted, as determined by hypergeometric distribution testing suggesting that EVs sampled from brain ISF with cOFM contains a brain-enriched miRNA signature (**Figure 3B**). Similarly, we compared circRNAs annotated from EV isolated from brain ISF samples with brain tissue expressing circRNAs established by Rybak-Wolf et al., who extensively profiled circRNAs in the mouse brain (Rybak-Wolf, Stottmeister et al. 2015). This comparison revealed a substantial overlap in annotated circRNAs in our ISF EV samples with those that showed high abundance in mouse brain (12,036 species) which was greater than predicted. Collectively, these comparisons support that EVs sampled from brain ISF with cOFM were enriched in brain expressing small ncRNAs. To reduce the dimensionality of the data and visualize differences in the types and abundance of annotated small ncRNA cargos from plasma and ISF EV samples we performed a Principal Component Analysis (PCA) analysis. We observed a clear separation between ISF and plasma EV samples, indicating compartment-specific small ncRNA abundance profiles (**Figure 3C**). To determine compartment-specific differences in small ncRNA abundance, we performed Deseq2 and identified 2,391 differentially abundant small ncRNA species in ISF EVs compared to plasma EVs (**Figure 3D, Supp. S1**). In ISF EVs, we identified a significant increase in 1,172 circRNA, 18 snoRNA, 135 snRNA, 38 miRNA, and 264 tRNA transcripts compared to plasma EVs, representing 9.5%, 3.8%, 56%, 6.3%, and 60% of the total annotated small ncRNA subtypes respectively, with snRNAs and tRNAs showing the greatest difference in abundance (**Figure 3E**). In contrast, plasma EVs showed a significant increase in 617 circRNA, 2 snoRNA, 7 snRNA, 110 miRNA, and 28 tRNA transcripts compared to ISF EVs, corresponding to 5%, 0.8%, 2.8%, 17%, and 6.3% of the total annotated small ncRNA subtypes, with miRNAs showing the highest difference in abundance (**Figure 3E**). These results highlight the unique small ncRNA profiles derived from ISF and plasma EVs. We focused on determining if the significantly increased small ncRNAs in ISF EVs compared to plasma EVs were also those that showed enriched brain expression utilizing the small ncRNA tissue expression atlas (Isakova, Fehlmann et al. 2020). In ISF EVs, we observed 38 miRNA transcripts that were upregulated with many showing enrichment in mouse brain tissue according to the atlas including: mir-218-5p, mir-223-3p, mir-323-3p, mir-344-3p, mir-434-3p and -5p, and mir-9-3p and -5p (**Figure 3F**, white). Additionally, several differentially abundant miRNAs higher in ISF EVs were previously determined to be expressed similarly in brain and other tissues, such as mir-149-5p, mir-219a-3p and mir-92b-3p (**Figure 3F**, light gray). We also identified novel miRNAs which were enriched in our ISF samples which were not previously identified as brain specific, like mir-5126-3p and mir-340-5p (**Figure 3F**, gray). In contrast, 110 miRNA transcripts were upregulated in plasma EVs compared to ISF EVs. Many of these miRNAs were enriched in peripheral tissues such as mir-133-3p (muscle enrichment), mir-122 (liver), and mir-375-3p (pancreas) (**Figure 3F**, yellow). We identified 1,172 significantly upregulated circRNA transcripts in EVs sampled from brain ISF compared to plasma, of which 1,109 (∼95%) overlapped with brain-expressed circRNA in the Rybak-Wolf dataset (**Figure 3G**, white) (Rybak-Wolf, Stottmeister et al. 2015). We observed only a few of the previously identified brain-expressing circRNAs had increased abundance in plasma EVs compared to ISF EVs (**Figure 3G**, yellow). The profiles of circRNAs identified as EV cargos from brain ISF strongly indicate that EVs sampled from the ISF using cOFM contain a brain-enriched signature. We utilized CircBase to identify the associated genes for the significantly increased circRNAs in ISF samples and determined that a large proportion were associated from genes that show high brain expression or encode for proteins that are important in many brain functions including: mmu_circ_0008013 (*Nrxn1*), mmu_circ_0008596 (*Rims1*), mmu_circ_0011123 (*Rims3*), mmu_circ_0012753 (*Gabra2*), and mmu_circ_0015971 (*Snap91*). In contrast, circRNA which were higher in abundance in plasma samples had associated genes that are higher expressing in the periphery, such as mmu_circ_0009539 (*Chd6*), mmu_circ_0013556 (*Sfxn5*), and mmu_circ_0001713 (*Cyld*). Gao et al. (2022) demonstrated tissue specific expression patterns of tRNAs across various mouse tissues, with brain tissue exhibiting markedly higher tRNA expression compared to other organs (Gao, Gallardo-Dodd and Kutter 2022). To assess if the EVs sampled from the ISF were enriched in a brain tRNA signature, we compared our dataset against the tRNA atlas reported by Gao et al. We identified 264 significantly upregulated tRNA transcripts in ISF EVs, with 43 overlapping with the enriched brain-expression (**Figure 3H**); supporting that EVs sampled from the ISF contain a brain-enriched small ncRNA signature. Next, we focused on Gene Ontology (GO) analysis for the identified predicted target genes of significantly increased miRNAs in ISF EVs using TargetScanMouse 8.0, aiming to uncover the potential functional roles of these miRNAs in regulating biological processes. We identified 2,333 predicted target genes and performed GO analysis with DAVID, categorizing the resulting GO terms into relevant biological processes (**Figure 3I**). GO analysis revealed strong enrichment in neuronal processes, including neuron generation, neuron development, neurogenesis, transmission of nerve impulse, brain development, and pathways critical for learning and memory. We postulate that EVs sampled from the ISF originate from the many different neural cell types highly organized in the brain including neurons, astrocytes and microglia. We wanted to determine if we could correlate the unique compartment specific small ncRNA signatures to published single-cell ATAC-seq data (Cusanovich, Hill et al. 2018) through comparing the expression scores of small ncRNAs predicted through ATAC-seq with the abundance of small ncRNAs in our brain ISF EV samples. The ultimate goal is to determine if the small ncRNA signature of the ISF EV samples were associated with different neural cell types. We found that several upregulated miRNAs in ISF EVs-including mir-124-1, mir-124-2, mir-124-3, mir-149-5p, mir-344-3p, and mir-409-3p were enriched from neuronal cell-type origin, while mir-136, mir-92b-3p, mir-9-3p, and mir-9-5p were both enriched in neurons and astrocytes (**Figure 3J**). Snord104 and Snord17 which were also increased in our ISF EV samples, demonstrate astrocyte and microglia cell-type origin, respectively (**Figure 3J**). These findings indicate that the increase abundance of small ncRNA transcripts in EVs isolated from ISF were associated with the major neural cellular types. Collectively, our comprehensive profiling of small ncRNAs across ISF and plasma EV samples showed that EVs sampled from the ISF were enriched in a brain small ncRNAs signature whose target genes encode for proteins important in brain functions. The small ncRNAs which were increased in the ISF EV samples were attributed to expression patterns in neurons, microglia, and astrocytes; further supporting that ISF EVs represents a distinct brain-derived EV population which is unique compared to EVs isolated from plasma.

**Figure 3.**
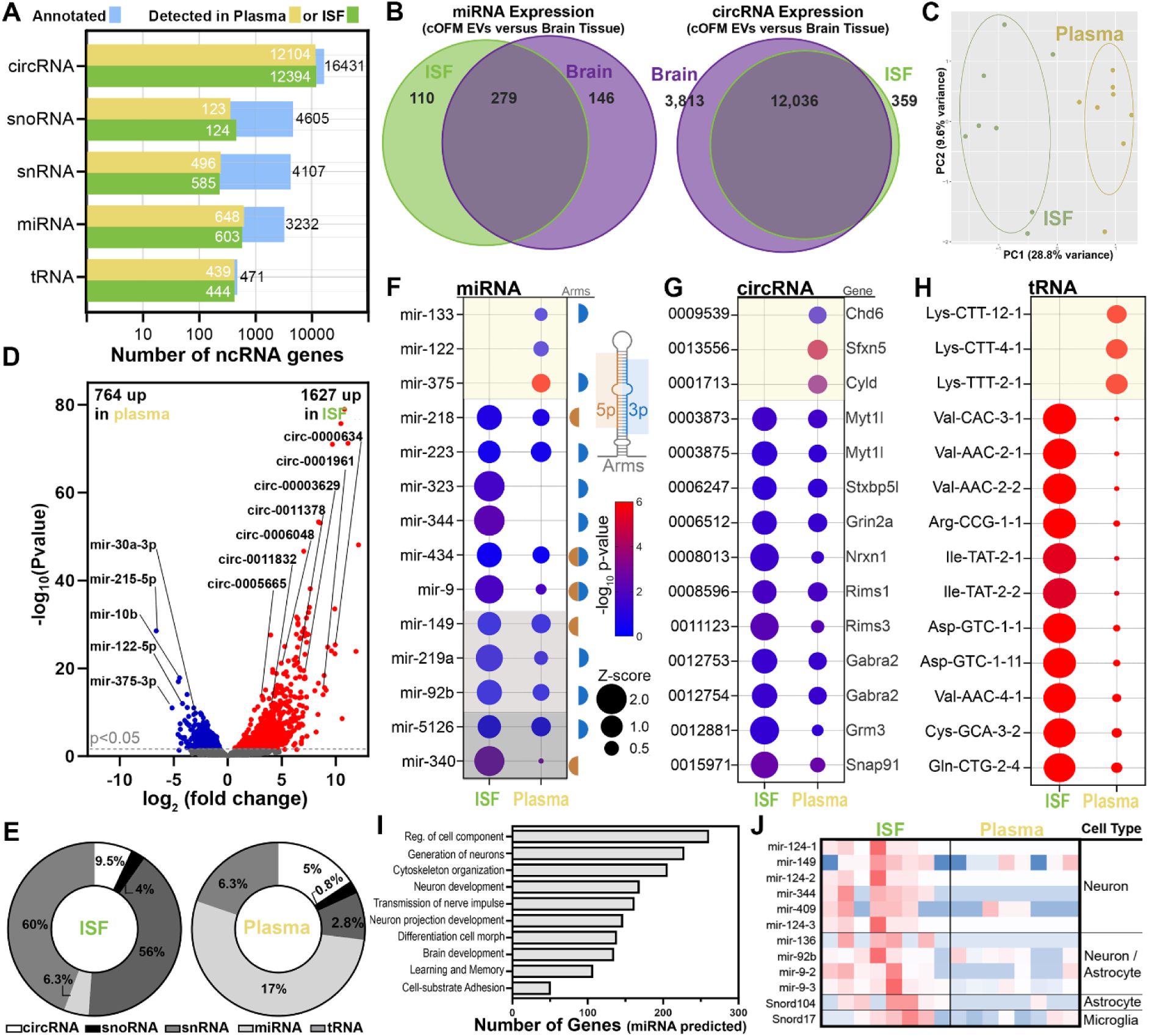
Unique EV populations were isolated from ISF and plasma based on small ncRNA enrichment profiles. (**A**) Bar plot of small ncRNAs annotated from ISF (green) and plasma EVs (yellow). Blue bars represent the total number of possible annotations in COMPSRA. (**B**) Venn diagrams representing the overlap between annotated small ncRNAs of the ISF EV samples and previous published data sets showing brain tissue enriched expression of miRNAs (**B**, Left) and circRNAs (**B**, Right). (**C**) PCA plot shows unique clustering of ISF and plasma EV samples based on small ncRNA abundance levels which illustrate distinct populations of EVs. (**D**) Volcano plot of differentially enriched small ncRNAs with 1627 significantly upregulated and 764 downregulated in EVs sampled from ISF by cOFM compared to plasma EVs (p<0.05). (**E**) Donut plots illustrate the distribution of significant upregulated small ncRNA transcripts in ISF EVs and plasma EVs. (**F**) Bubble plots showing a summary of the identified brain enriched miRNA transcripts (Isakova, Fehlmann et al. 2020) that were upregulated in ISF EV samples compared with plasma EVs. Color denotes: miRNAs what were upregulated in plasma EV samples and show enriched expression in peripheral tissues (light yellow), miRNAs what were upregulated in ISF EV samples and display enriched expression in brain tissue (white), miRNAs what were upregulated in ISF EV samples and are expressed in brain tissue (light gray), and miRNAs what were upregulated in ISF EV samples and previous have not shown to have brain tissue expression (deep gray). “Arm” column denotes whether 5p-, 3p-, or both arms of the miRNA species shows significant differential enrichment. (**G**) Bubble plots showing a sample of the circRNAs with enriched brain tissue expression (Rybak-Wolf, Stottmeister et al. 2015) and identified as significantly increased in abundance in the ISF EV samples compared to plasma EVs. Color denotes: circRNA what were upregulated in plasma EV samples and show enriched expression in peripheral tissues (light yellow), circRNA what were upregulated in ISF EV samples and display enriched expression in brain tissue (white). (**H**) tRNAs which were significantly increased in ISF EV samples compared to plasma EVs and identified as brain enriched previously (Gao, Gallardo-Dodd and Kutter 2022). Color denotes: tRNA what were upregulated in plasma EV samples and show enriched expression in peripheral tissues (light yellow), tRNA what were upregulated in ISF EV samples and display enriched expression in brain tissue (white). For all bubble plot, bubble color represents statistical significance (-log₁₀ p-value) and bubble size corresponds to normalized gene expression. (**I**) Bar plot showing top Gene Ontology (GO) categories of miRBase predicted target genes from miRNAs that were upregulated in ISF EVs compared to plasma EVs samples. Bar size represent the number of genes and all terms are significant at p<0.05 (**J**) Heatmaps display cell-type origin specific expression patterns of significantly differentially enriched small ncRNAs identified in EVs from ISF and plasma compartments, based on (Cusanovich, Hill et al. 2018). Circular RNA; snoRNA: Small nucleolar RNA; snRNA: Small nuclear RNA; miRNA: MicroRNA; and tRNA: Transfer RNA. n = 8 mice/group (4 WT and 4 APP/PS1 with equal sex distribution).

### 3.4 Distinct small ncRNA signatures in EVs sampled from brain ISF of AD model mice

Once we demonstrated that EVs sampled from brain ISF were enriched in a neural small ncRNA signature, we next wanted to determine whether these EVs could be used to discriminate changes in the small ncRNA abundance associated with neurodegenerative diseases, such as AD. To address this, we compared the small ncRNA profiles of EVs isolated from brain ISF, brain tissue, and plasma of WT and APP/PS1 mice. We identified 349 significant differentially enriched small ncRNAs transcripts in APP/PS1 mice compared to controls from brain ISF isolated EVs (**Figure 4A, Supp. S2**). Furthermore, we identified 807 differentially abundant small ncRNA transcripts in brain tissue EVs (**Figure 4B, Supp. S2**), and 445 differentially abundant small ncRNA transcripts in plasma EVs when comparing APP/PS1 to WT mice (**Figure 4C, Supp. S2**). These findings indicate that each compartment exhibits distinct alterations in EV-associated small ncRNA profiles associated with AD pathology. Donut plot revealed compartment-specific differences in small ncRNA transcripts abundance within EVs isolated from brain ISF, brain tissue, and plasma of APP/PS1 mice compared to WT controls (**Figure 4D**). In EVs sampled from brain ISF by cOFM, we identified significant alterations in 305 circRNAs, 4 snoRNAs, 18 snRNAs, 19 miRNAs, and 3 tRNAs, representing 2.5%, 3.2%, 3.0%, 3.1%, and 0.7% of the total annotated transcripts in each respective class. EVs isolated from brain tissue had significant changes in 479 circRNAs, 7 snoRNAs, 41 snRNAs, 174 miRNAs, and 136 tRNAs, representing 3.2%, 2.7%, 2.6%, 9.5%, and 30% of the total annotated transcripts in each class. Plasma EVs exhibited significant differences in 273 circRNAs, 11 snoRNAs, 26 snRNAs, 68 miRNAs, and 67 tRNAs, corresponding to 2.3%, 10%, 5.5%, 10.7%, and 15% of the total annotated transcripts in each class, respectively). Collectively, these data highlight distinct and compartment-specific small ncRNA signatures in APP/PS1 mice when compared to WT controls.

**Figure 4.**
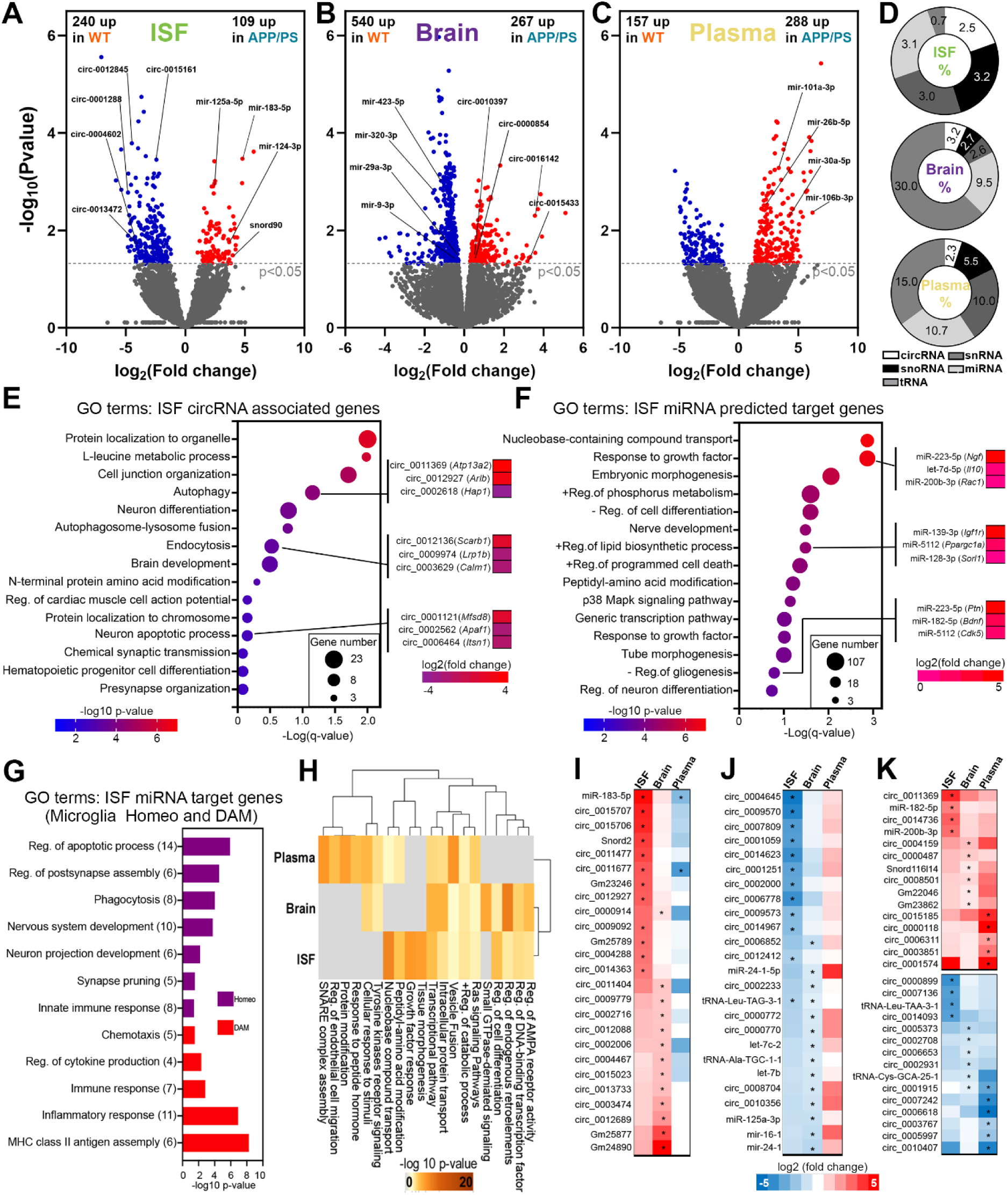
Distinct small ncRNA signatures in EVs sampled from brain ISF of AD model mice. Volcano plot of differentially enriched small ncRNAs transcripts between APP/PS1 and WT mice of isolated EVs from (**A**) ISF (109 up- and 240 downregulated in APP/PS1), (**B**) Brain (267 up- and 540 downregulated in APP/PS1), and (**C**) Plasma (288 up- and 157 downregulated in APP/PS1). (**D**) Donut plots illustrate the percent distribution of significant differential abundant small ncRNA transcripts in EVs when comparing APP/PS1 and WT mice from brain ISF, brain tissue, and plasma EV samples. Bubble plots depicting significantly enriched GO terms derived from predicted associated genes of differentially enriched circRNAs (**E**) and miRNAs (**F**) identified in ISF EVs from APP/PS1 compared to WT mice. Bubble size indicates the number of genes within each GO term, and color intensity corresponds to statistical significance. Accompanying heatmaps demonstrate the expression profiles (log2 fold change) of selected genes associated transcripts from representative GO categories, emphasizing differential regulation between APP/PS1 and WT mice. (**G**). miRNA predicted target genes classified as either homeostatic (Homeo, pink bars) or disease-associated microglia (DAM, red bars) based on comparison with published single-cell RNA sequencing datasets (Lu, Saibro-Girardi et al. 2023). GO analysis highlights biological processes distinctly associated with these microglial phenotypes. (**H**) Heatmap and dendrograms show commonality between GO terms enriched from with the gene targets of different abundant miRNAs in ISF, brain tissue, and plasma compartments of APP/PS1 and WT mice. (**I-J**) Heatmaps illustrating expression (log2 fold change) of selected small ncRNA species with similar enrichment in EVs from ISF and brain compared to plasma of APP/PS1 versus WT mice. Statistical significance (*p < 0.05) indicated by asterisks. N=8 mice/group with equal male female ratio (APP=4 and WT=4).

In the present study, we focused on small ncRNAs derived from brain ISF EVs of APP/PS1 compared to WT mice to uncover key regulatory networks and biological processes influenced by altered small ncRNA EV cargos, providing insights into mechanisms associated with AD pathology. In EVs isolated from brain ISF, we identified 305 significantly different circRNA transcripts in APP/PS1 mice compared to WT controls, highlighting substantial changes in the circRNA cargos associated with AD pathology. To gain deeper insights into the biological significance of these circRNA cargos, we performed target gene prediction using CircBase and identifying 202 putative associated genes. The list of genes predicted by CircBase were then analyzed with Metascape to determine associated biological processes. Notably enriched GO terms included autophagy, endocytosis, brain development, neuron apoptosis, and presynaptic organization, suggesting important regulatory roles for circRNA-targeted pathways in the modulation of neuronal health and function in AD model mice (**Figure 4E**). Autophagy dysfunction is a known contributor to AD pathology due to impaired clearance of misfolded proteins such as Aβ and hyperphosphorylated tau, leading to their toxic accumulation and subsequent disruption of cellular homeostasis and synaptic function (Barmaki, Nourazarian and Khaki-Khatibi 2023). In ISF EVs, the circRNA associated genes related to autophagy included, *Atp13a2* (circ_0011369), a gene linked to lysosomal dysfunction during neurodegeneration leading to impaired acidification and autophagosome clearance (Dehay, Martinez-Vicente et al. 2013); *Arl8b* (circ_0012927), which is involved in lysosomal transport and accumulates on axonal lysosomal membranes near Aβ plaques (Boeddrich, Haenig et al. 2023); and *Hap1* (circ_0002618), a neuronal protein important in trafficking of APP and shown to modulate levels of amyloid plaque (Yang, Yang et al. 2012). This highlights that APP/PS1 mice exhibit disease-related changes in circRNA cargos which could be associated with autophagy, critical for in AD pathology. Furthermore, dysregulation of endocytic pathways can enhance the production of Aβ peptides through increased internalization of APP and its subsequent cleavage (Cirrito, Kang et al. 2008). In the endocytosis GO term we identified circRNA associated genes, *Scarb1* (circ_0012136), *Lrp1b* (circ_0009974), and *Calm1* (circ_0003629), all which encode for proteins important in modulating amyloid pathology (Cam, Zerbinatti et al. 2004, Wu, McNeil et al. 2009, Thanopoulou, Fragkouli et al. 2010). Here we highlight the potential of cOFM to collect brain ISF EVs in APP/PS1 mice which have significantly different circRNA cargos than WT controls. The differently enriched circRNA were associated with genes that are part of biological processes important in amyloid pathology, revealing the potential involvement of these brain ISF EV cargos in key cellular processes and pathways associated with neurodegeneration.

Similarly, in EVs isolated from brain ISF, we identified 19 significantly enriched miRNA transcripts in APP/PS1 mice compared to WT controls, highlighting changes in the miRNA cargos associated with AD pathology. Using TargetScan, we predicted 1,068 target genes using (cut off value at TCS>-0.5) and performed GO enrichment analysis using Metascape. We identified enrichment in key biological processes related to response to growth factor stimuli, nerve development, positive regulation of lipid biosynthetic process, regulation of neuron differentiation and neurodegeneration (**Figure 4F**). Notably, many of these pathways are closely linked to neurodegenerative pathology, suggesting that the altered miRNA cargo in ISF EVs may contribute to disease progression by modulating gene expression. For example, growth factor signaling is essential for neuronal survival, synaptic plasticity, and cognitive integrity-all of which are disrupted in AD. Among the predicted target genes associated with the growth factor response included *Ngf* (mir-223-5p), *Il-10* (let-7d-5p) and, *Rac1* (mir-200b-3p). Not only do these genes encode proteins important in growth factor response in neurons but they also modulate the role in microglia. Both NGF and IL-10 play anti-inflammatory roles in the brain and can influence microglial function and reduce neuroinflammation (Lobo-Silva, Carriche et al. 2016, Rizzi, Tiberi et al. 2018). Where coordination of *Rac1* plays a key role in determining phagocytic potential of microglia (Kim, Kim et al. 2017). With increased abundance of these miRNAs in the ISF EVs of APP/PS1 mice this could drive decreased expression of these key genes leading to an increased inflammatory state and decreased microglia phagocytosis. Interestingly, we also saw enrichment of miRNA in the ISF EVs of APP/PS1 mice which regulate genes (*Ptn, Bdnf, Cdk5*) important in gliogenesis; highlighting the potential of these miRNA EV cargos to modulate glial functions important in responding to amyloid pathology. AD is increasingly recognized as being intertwined with the dysregulation of lipid metabolism (Fitz, Wang et al. 2021). Interestingly, we see enrichment of miRNAs in the ISF EVs of APP/PS1 mice compared to WT controls with targets genes that express proteins important in the regulation of lipid biosynthesis and other metabolic functions (*Igf1r, Ppargc1a, Sorl1*). This data suggests that brain ISF EVs is a unique liquid biopsy which can be used to monitor changes in cargos that might modulate AD related pathologies.

With increased understanding of the significant multifaceted role that microglia play in AD and the number of biological processes and inflammatory related genes which we determined were targets of the miRNA cargos in ISF EVs; we compared these miRNA predicted gene targets to our published single cell RNAseq of differential expressed genes in both homeostatic (homeo) and disease associated microglia (DAM) populations (Lu, Saibro-Girardi et al. 2023). To explore microglia-specific biological alterations, we compared the TargetScan-predicted genes from ISF EV miRNAs with the single-cell data. This analysis identified 155 DAM-associated miRNA target genes enriched in processes such as regulation of apoptosis, postsynapse assembly, phagocytosis, nervous system development, synapse pruning, and innate immune response (**Figure 4G**). In contrast, 433 Homo-associated miRNA target genes were linked to MHC class II antigen presentation, inflammatory and immune responses, cytokine regulation, and chemotaxis (**Figure 4G**). These findings highlight the complex nature by which brain EV miRNA cargos can influence microglial states in AD and the usefulness of cOFM for monitoring these changes during different stages of AD progression. Taken together, our results demonstrate that ISF-derived EVs carry a distinct small ncRNA profile enriched in neurodegeneration-related pathways.

To determine whether ISF EVs reflect brain-specific molecular alterations associated with AD, we compared common ncRNA transcripts, predicted associated genes, and GO terms across ISF, brain, and plasma EVs in APP versus WT mice (**Figure 4H-K**). When analyzing the GO terms associated with the TargetScan gene targets of the differential enriched miRNAs in APP/PS1 mice compared to WT mice of EVs isolated from ISF, brain tissue and plasma the dendrogram clustering demonstrates higher similarity between brain and ISF compartments compared to plasma (**Figure 4H**). These include genes important in biological processes which were unique to brain tissue and ISF compartments, such as cell differentiation, host-immunity responses, and AMPA receptor activity which is important in synaptic plasticity (Chater and Goda 2014). Where the plasma EV compartment had miRNA target genes enriched with unique biological processes associated with peptide hormones, endothelial cell migration and SNARE complex. We also observed several enriched small ncRNAs when comparing APP/PS1 mice to WT in both ISF and brain tissue EVs that were downregulated in plasma. (**Figure 4I**). Conversely, several significantly downregulated small ncRNAs transcripts in ISF and brain EVs were found to be upregulated in plasma EVs (**Figure 4J**). These opposing trends emphasize that plasma EVs may not reliably represent brain-derived changes due to peripheral influences or additional systemic signal. In contrast, ISF and brain EVs exhibited concordant expression patterns, suggesting that ISF EVs closely reflect brain molecular dynamics and may serve as a more accurate and accessible liquid biopsy for monitoring CNS-specific pathological processes in AD. There were few transcripts that exhibited a consistent direction of fold change across ISF, brain, and plasma EVs in APP/PS1 versus WT mice. (**Figure 4K**). Taken together, the predicted target genes GO enrichment, along with the targeting of key microglial populations, close molecular similarity between ISF and brain EVs, suggests that ISF EVs play an active role in modulating AD related pathologies. Thus, our findings demonstrate the usefulness of cOFM for monitoring pathological changes in ISF EVs and provide important insight into how ISF EVs may contribute to neurodegenerative disease like AD progression and highlight their potential as early indicators of disease-related changes.

## 4. Discussion

Microdialysis has long been established as a technique for sampling proteins of the brain ISF and monitoring dynamic biochemical changes *in-vivo*. In the context of neurodegenerative diseases and brain injury, microdialysis has been widely applied to measure soluble proteins, metabolites, and neuropeptides over time. For example, microdialysis in AD models have shown that Aβ levels fluctuate in response to neuronal activity and that therapies targeting Aβ production can modulate these levels in real time (Magnoni, Esparza et al. 2012, Fitz, Cronican et al. 2013, Shahnur, Nakano et al. 2021). Despite its established use in measuring single protein biomarkers, the application of microdialysis for capturing complex molecular carriers such as EVs has remained largely unexplored.

Our study demonstrates for the first time that ISF EVs can be isolated using OFM sampling and the small ncRNA cargos molecularly profiled. The OFM sampling is not restricted by a membrane which might otherwise limit the efficiency of recovery and the populations of EVs which could be samples (Pait, Kaye et al. 2024). We applied cOFM in WT and APP/PS1 mice to sample brain ISF-derived EVs which had similar characteristics as EVs isolated from plasma or brain tissue. Unlike brain tissue, which is a solid compartment requiring invasive procedures for EV isolation, both ISF and plasma are liquid compartments that can be accessed with minimal tissue disruption. This makes them more suitable for studying EVs in a physiologically relevant and dynamic manner. ISF EVs, being directly shed from brain-resident cells, offer brain-specific insights, while plasma EVs provide a minimally invasive route to monitor systemic and brain-derived changes. While the analysis of the small ncRNA cargos identified in EVs sampled from brain ISF and plasma EVs had similar number of annotated small RNA species there was a significant difference in the abundance of these small ncRNAs between the two compartments (**Figure 3**). We found that the majority of miRNA (204 of 247) and circRNA (12,036 of 12,395) transcripts annotated in EVs sampled from the brain ISF by cOFM were highly enriched in whole brain tissue, indicating that ISF EVs carry a molecular signature highly reflective of brain origin. Differential enrichment analysis further revealed the differences between the two compartments, with 1,627 small ncRNAs transcripts significantly increased in brain ISF EVs and 764 significantly increased in plasma EVs. Notably, brain ISF EVs exhibited marked enrichment in circRNA transcripts, snRNA transcripts, and tRNA transcripts while plasma EVs showed greater enrichment of miRNA transcripts. This supports other research demonstrating that circRNAs are generally more abundant and diverse in the mammalian brain than in other tissues, enriched at the synapses, conserved across species, and linked to neuronal development and aging (Westholm, Miura et al. 2014, Rybak-Wolf, Stottmeister et al. 2015, You, Vlatkovic et al. 2015). Comparison of the miRNAs enriched in the EVs sampled from ISF with published tissue atlas of small ncRNA expression (Isakova, Fehlmann et al. 2020) demonstrated that most of these miRNAs also had higher expression in the brain compared to other tissues, including mir-9-3p, miR-218-5p, mir-223-3p, and mir-323-3p. mir-9-3p is a key regulator of synaptic plasticity and memory formation, influencing neuronal connectivity and cognitive processes (Sim, Lim et al. 2016). mir-218-5p has been shown to protect dopaminergic neurons (Ma, Zhang et al. 2021) and important in neural responses to chronic stress and depression (Yoshino, Roy and Dwivedi 2022). While mir-223-3p plays a neuroprotective role in modulating neuroinflammation (Morquette, Juzwik et al. 2019). Similarly, the abundant of circRNA in ISF EVs are associated with genes that are highly enriched in the brain. For example, circ_0008013 (derived from *Nrxn1*), members of the neurexin family, are critical regulators of synapse formation, maintenance, and signaling (Sudhof 2008). Similarly, circ_0008596 (derived from *Rims1*) and circ_0011123 (derived from *Rims3*) are essential for neurotransmitter release and synaptic plasticity, fundamental processes underpinning proper brain function (Wu, Fan et al. 2023). In addition, circ_00 12753 (derived from *Gabra2*) originates from a gene encoding the alpha-2 subunit of the GABA-A receptor, a key modulator of neuronal excitability (Gonzalez-Nunez 2015), while circ_0015971 (derived from *Snap91*) corresponds to a synapse-associated protein highly enriched in brain (Ishikawa, Nagase et al. 1998).

Beyond establishing brain enrichment of EVs and their small ncRNA signature, our functional analyses provide additional insight into the potential roles of these ISF EV cargos. Gene ontology analysis performed with the gene targets of the enriched miRNAs in ISF EV revealed strong association in pathways central to neuronal development, synapse organization, neurotransmission, and neurogenesis-biological processes critical for brain homeostasis and plasticity. Importantly, enriched miRNAs in ISF EVs isolated in our study mapped predominantly to neuronal, astrocytic, and microglial expression patterns (Cusanovich, Hill et al. 2018), further supporting the hypothesis that multiple brain cell types contribute to the EV pool within the brain ISF. Together, these findings align with prior studies implicating these miRNAs, circRNAs in key CNS pathways, suggesting that ISF EVs could serve as functional mediators of intercellular communication within the brain and the cOFM would be a valuable technique to monitor these changes *in vivo*.

In contrast to the isolated brain ISF EVs, our analysis showed that plasma EVs were enriched in peripheral tissue-associated miRNAs, including mir-122, mir-133-3p, and mir-375-3p, each reflecting specific organ origins and biological roles. mir-122 is predominantly expressed in the liver, where it regulates lipid metabolism, cholesterol homeostasis, and plays a critical role in hepatocellular carcinoma development and liver injury responses (Tsai, Hsu et al. 2012). mir-133-3p is highly expressed in cardiac and skeletal muscle, contributing to muscle differentiation, regeneration, and protection against cardiac hypertrophy (Horak, Novak and Bienertova-Vasku 2016, Crocco, Montesanto et al. 2024). The enrichment of these miRNAs in plasma EVs highlights the contribution of systemic tissues to peripheral circulating EV cargos, contrasting with the brain-enriched miRNA profile observed in the sampled ISF EVs. This further illustrates that the dynamic secretion of EVs from different peripheral tissues and their unique small ncRNA expression patterns makes it difficult to use plasma as a source for monitoring changes in EV cargos associated with the brain development, aging or neurodegeneration. Thus, using small ncRNA profiling, we show that EVs isolated from ISF carry a highly brain-enriched molecular cargo, distinct from plasma EVs, and each compartment harbors a distinct small ncRNA signature, reflecting compartment-specific molecular compositions.

EVs play critical roles in brain, carrying cargos that include proteins, lipids, and small RNAs that reflect not only the physiological state but also neurodegenerative processes. The EV cargos could serve as a biomarker of brain health and contribute to normal homeostatic functions or pathology. Notably, several studies have reported disease-associated alterations in EVs cargo relevant to AD and other neurodegenerative conditions. For example, increased levels of phosphorylated tau (p-tau) in brain- and CSF derived EVs have been reported in AD patients and animal models, suggesting that EVs may contribute to the spread of tau pathology between neurons and serve as sensitive biomarkers of disease progression (Saman, Kim et al. 2012, Polanco, Scicluna et al. 2016). Furthermore, studies have shown that microtubule-associated protein tau circular RNAs can be translated and facilitate tau tangle formation (Welden, Margvelani et al. 2022), which could be a unique mechanism by which EV cargos can facilitate the spread of tau pathologies. With the utilization of cOFM there is the potential to better define changes in other cargos such as lipids, metabolites and proteins associated with brain health and disease. This further characterization of brain ISF EVs and their cargos could be crucial for clearly defining biomarkers of neurodegeneration.

We demonstrate that cOFM sampling of the ISF EVs is a powerful approach to capture brain-specific molecular changes that are otherwise diluted or masked in peripheral biofluids such as plasma. A major advantage of ISF sampling lies in its unique ability to track molecular changes in the brain over time, achieving longitudinal access to the extracellular environment of the brain with repeated sampling from the same mouse at multiple time points. This temporal flexibility contrasts with brain tissue or even plasma sampling, which typically relies on single or few time-points and provides only static molecular snapshots. As neurodegenerative diseases such as AD are progressive and temporally dynamic, the ability to monitor EV cargo in real-time offers a significant methodological and conceptual advance. ISF-derived EVs, therefore, provide an unparalleled window into the ongoing importance of EV in molecular changes within the brain, allowing us to capture early and progressive molecular events during disease progression and how therapeutics could modulate these changes. Thus, ISF EVs serves as a valuable and dynamic source for monitoring EVs changes during neurodegeneration. Considering these findings, we next explored how the brain-enriched small ncRNA cargos of ISF EVs are altered in the context of neurodegenerative disease, focusing on AD.

Differential enrichment analysis revealed substantial changes in the small ncRNA abundance in ISF EV of APP/PS1 mice compared to WT with 109 up- and 240 downregulated species indicating unique small ncRNA signatures associated with amyloid pathology (**Figure 4A**). Specifically, ISF EVs exhibited significant changes in 305 circRNAs, 19 miRNAs, and smaller yet biologically relevant numbers of snoRNAs, snRNAs, and tRNAs (**Figure 4D**). The relatively balanced distribution across small ncRNA classes suggests that ISF EVs may capture early or localized alterations in neuronal functions with circRNAs, highly enriched and stable in the brain and implicated in neurodegenerative diseases such as AD (Rybak-Wolf, Stottmeister et al. 2015, Dube, Del-Aguila et al. 2019, Zhang and Bian 2021). In this study, GO enrichment analysis of circRNA predicted target genes from ISF EVs further revealed their functional involvement in biological pathways highly relevant to neurodegeneration including autophagy, endocytosis, presynaptic organization, and neuron apoptosis (**Figure 4E**). For example, several downregulated circRNA circ_0012845, circ_0001288, circ_0004602, and circ_0013472 (**Figure 4A**) identified in ISF EVs from APP/PS1 mice were predicted to target genes with established roles in AD pathogenesis. For instance, circ0012845 is associated with *Dpp6*, a gene involved in synaptic plasticity and linked to AD-related cognitive decline (Cacace, Heeman et al. 2019). circ0001288 with *Ubr4*, an E3 ubiquitin ligase implicated in protein quality control and neuroprotection (Haakonsen, Heider et al. 2024) and circ0004602 with *Ranbp9*, a scaffold protein that enhances amyloid precursor protein processing and Aβ production (Lakshmana, Chung et al. 2010). The downregulation of these circRNAs may reflect impaired regulatory control over key AD-related pathways, aligning with prior studies implicating circRNA dysregulation in AD progression. Similarly, ISF EV-enriched miRNAs in APP/PS1 mice compared to WT were found to target genes related to many functional GO terms such as response to growth factor stimuli, nerve development, lipid biosynthesis regulation, regulation of neuron and glia differentiation (**Figure 4H**). miRNAs are known to regulate post-transcriptional gene expression and have been implicated in AD-related processes including synaptic dysfunction, neuroinflammation, and impaired protein trafficking (Lau, Bossers et al. 2013, Rajgor and Hanley 2016). In line with this, several miRNAs such as mir-125a-5p, mir-183-5p, mir-124-3p (**Figure 4A**) upregulated in ISF EVs from APP/PS1 mice are associated with AD pathogenesis. For example, mir-125a-5p found in serum EVs was identified as a potential biomarker in late-onset AD (de Lourdes Signorini-Souza, Tureck et al. 2024) and in the current study we were able to capture similar patterns in the sampled EVs from brain ISF. mir-183-5p reduces neuronal apoptosis (Zhu, Zhou et al. 2020); while mir-124-3p decreased p25 formation (Zhou, Deng et al. 2019), both modulating neuronal protection. These findings highlight the potential of ISF EV-derived miRNAs in AD to not only serve as a biomarker but high significant biological relevance to AD related phenotypes.

Comparison of the miRNA target genes with our single-cell RNA-seq gene expression dataset (Lu, Saibro-Girardi et al. 2023) further demonstrated that miRNA carried by ISF EVs can target genes that express proteins important in regulation of both homeostatic and disease-associated microglial states (Homeo and DAM) (**Figure 4H**). DAM-associated targets were linked to synaptic pruning, phagocytosis, apoptosis, and innate immune pathways processes central to neurodegenerative progression. After the initial characterization of the DAM microglia (Keren-Shaul, Spinrad et al. 2017) in AD mouse models there has been an increased focus on determining how these microglia states are regulated. Here we present the potential of small ncRNAs delivered by brain ISF EVs as a potential regulator of microglial functions. Collectively, these findings suggest that ISF EV-derived small ncRNAs not only coordinate core developmental pathways but also contribute to neuroinflammatory processes and neurodegeneration associated with AD.

Importantly, the EVs sampled from the ISF with cOFM had a similar enrichment profile in APP/PS1 mice compared to EVs directly isolated from the brain tissue of the same mouse at the end of the experiment (**Figure 4G-J**). We observed several similar associated biological processes, such as cellular differentiation, inflammatory processes, regulation of AMPA signaling and transcriptional regulations, when examining the gene targets of the enriched miRNAs in EVs from both compartments of APP/PS1 mice compared to controls. This was highlighted by similar enrichment profiles of the small ncRNAs in brain and ISF samples compared to plasma. A subset of ncRNAs exhibited consistent directional changes across all three compartments, such as circ_0011369 and miR-182-5p (upregulated), and circ_0000899 and tRNA-Leu-TAA-3-1 (downregulated), indicating their potential as stable molecular indicators of AD-associated pathology (**Figure 4K**). Included in this enrichment signature in APP/PS1 mice across all three compartments was a species belonging to the SNORD116 family, which we previously demonstrated was increased in plasma EVs isolated from AD patients (Fitz, Wang et al. 2021). This not only emphasizes the similarities between EVs isolated from brain tissue and sampled from the ISF, demonstrating that cOFM samples EV that are a reliable proxy for brain molecular signaling, but also shows the potential usefulness of cOFM for sampling over dynamic windows during the pathological progression which is a limitation of EV isolated from brain tissue.

cOFM represents a transformative approach for accessing ISF EVs directly from the brain parenchyma, enabling real-time monitoring of neurobiological processes during development and in both acute and chronic disease contexts. Unlike traditional sampling methods, cOFM allows for continuous and dynamic collection of intact nanoscale particles including EVs without the limitations of molecular size cutoff, making it particularly valuable for studying the complex pathophysiology of neurodegenerative diseases. For example, in traumatic brain injury (TBI) model, cOFM-sampled ISF EVs could serve as sensitive indicators of ongoing neuroinflammatory responses. Studies show that EVs secreted by activated microglia, astrocytes, and infiltrating immune cells, carry a wide array of bioactive molecules such as cytokines, chemokines, miRNAs, and damage-associated molecular patterns that drives secondary injury and inflammatory cascade (Liu, Zhang et al. 2022). In addition to local release, their systemic dissemination detectable in serum and cerebrospinal fluid suggests that ISF EVs can be detected beyond the injury site (Hazelton, Yates et al. 2018, Manek, Moghieb et al. 2018). Thus, cOFM-sampled ISF EVs can reflect ongoing pathological changes in the brain through accessible biofluids makes them promising candidates for non-invasive biomarkers and potential modulators of therapeutic response in TBI. Beyond acute injury, cOFM can be uniquely suited to elucidate EV-mediated mechanisms underlying chronic neurodegeneration. In tauopathies, brain ISF EVs sampled by cOFM can encapsulate misfolded, hyperphosphorylated tau species that could facilitate their intercellular transfer and pathological seeding (Leroux, Perbet et al. 2022). This EV-mediated propagation could contribute to the spread of tau across anatomically connected brain regions. Similarly, in synucleinopathies, such as Parkinson disease, EVs can carry monomeric and aggregated α-synuclein, promoting neuronal dysfunction and glial activation (Shippey, Campbell et al. 2022, Li, Huang et al. 2024). Here we highlighted differences in the small ncRNA cargos associated with amyloid pathology, but other cargos could modulate Aβ production, aggregation, clearance, and associated neuroinflammation (Petit, Fernandez et al. 2022). The capacity of cOFM for continuous monitoring of brain ISF EVs enables the mapping of early pathogenic events and disease progression trajectories in the brain.

Here we show proof-of-concept for sampling ISF EV by cOFM followed by small ncRNA profiling to determine changes in AD model mice. Furthermore, translating these findings other neurodegenerative models and during different stages of the disease progression could provide a novel platform for biomarker discovery and therapeutic monitoring. While CSF and plasma EVs are already being explored as AD biomarkers, ISF EVs offer the advantage of direct access to brain-specific changes with minimal peripheral dilution and the ability to sample in the same mouse over time. Here we define a unique small ncRNA signature in ISF EV which has the potential to target important biological process related to amyloid pathology and modulate microglial gene networks, emphasizing the potential of ISF EVs to modulate of disease progression and neuroinflammatory pathways. These EV associated small ncRNAs particularly could serve as accessible biomarkers for early-stage AD diagnosis and progression monitoring due to their presence in biofluids and resistance to degradation. By offering a real-time snapshot of the brain’s molecular landscape, ISF EVs hold promise for advancing both the understanding and clinical monitoring of neurodegenerative diseases and for guiding the development of small ncRNA-based biomarkers or EV directed therapies.

**Supplemental Table 1**

Differentially enriched small ncRNAs in cOFM compared to plasma.

**Supplemental Table 2**

Differentially enriched small ncRNAs in AD model mice compared to WT mice from cOFM, brain and plasma.

## Availability of data and materials

Raw data was submitted to GEO (https://www.ncbi.nlm.nih.gov/geo/) and will be publicly available as of the date of publication. This paper does not report the original code. Any additional information required to reanalyze the data reported in this paper is available from the corresponding author upon request.

## Funding

This work was funded by the National Institute on Aging, National Institutes of Health, USA (AG075069-NFF; AG077636-RK / IL; AG075992-IL / RK).

## Authors’ contributions

Conceptualization, N.F.F; Methodology, N.F.F. M.D.A; Investigation, M.D.A., S.A., M.A.O, N.F.F.; Formal analysis, M.D.A., S.A., M.A.O, N.F.F.; Visualization, M.D.A., S.A., M.A.O, N.F.F.; Writing – Original draft, M.D.A., N.F.F.; Writing – Review & Editing, R.K., I.L.,; Supervision, R.K., I.L., N.F.F.; Project administration, R.K., I.L., N.F.,F.; Resources, R.K., I.L., N.F.F.; Funding acquisition R.K., I.L. N.F.F.

## Ethics declarations

Ethics approval and consent to participate

All animal procedures were performed according to the Guide for the Care and Use of Laboratory Animals from the United States Department of Health and Human Services, and were approved by the University of Pittsburgh Institutional Animal Care and Use Committee.

## Consent for publication

Not applicable.

## Competing interests

The authors declare no conflict of interest.

## Notes

### Competing Interest Statement

The authors have declared no competing interest.

